# Genomic insights into the genus *Pantoea*: biotechnological potential and lifestyle diversity

**DOI:** 10.1101/2025.10.01.679771

**Authors:** Felipe F. Rimes-Casais, Francisnei Pedrosa-Silva, Thiago Motta Venancio

## Abstract

*Pantoea* is a genus of Gram-negative bacteria isolated from diverse environments. Over time, it has drawn considerable attention for its potential to promote plant growth. However, its biotechnological application is complicated by high genomic plasticity, which underlies both its beneficial traits and its ability to cause disease in a wide range of plants, as well as occasional opportunistic infections in humans, raising biosafety concerns. In this study, we conducted a comparative genomic analysis of all publicly available *Pantoea* genomes. Our goals were to refine taxonomic classifications and to identify genes linked to biotechnological potential, virulence, and antibiotic resistance, thereby clarifying lifestyle strategies within the genus. We found that plant growth-promoting genes are widely conserved, particularly those involved in phosphate solubilization, phytohormone biosynthesis, and siderophore production. In contrast, traits such as nitrogen fixation and ACC deaminase activity were restricted to specific species. The resistome analysis revealed intrinsic resistance mechanisms conserved across the genus, primarily involving diverse efflux pump families and β-lactamases conferring resistance to cephalosporins. In parallel, the pan-GWAS highlighted lifestyle-defining genetic markers, including the *hrp/hrc* genes encoding type III secretion system components, *pepM* (phosphoenolpyruvate mutase) associated with the production of a phytotoxin, and *ibeB*, an invasin linked to clinical infections. Together, our findings underscore both the biotechnological potential of *Pantoea* and the importance of genetic markers for distinguishing beneficial from pathogenic lifestyles, supporting the safe application of selected strains in biotechnology.

## Introduction

*Pantoea* is a genus of Gram-negative, motile, rod-shaped bacteria belonging to the family Erwiniaceae. The genus name, meaning “of all types and sources”, reflects its broad ecological distribution and diversity. Members of *Pantoea* are frequently isolated from plants, where they are well-known colonizers with strong competitive ability and remarkable adaptability to diverse hosts (Crosby et al., 2023). Beyond plants, these bacteria have also been recovered from soil, water, in commensal interactions with insects, and in urban environments (Walterson; Stavrinides, 2015), underscoring their versatile and wide-ranging ecological lifestyle.

Over the past decades, *Pantoea* has attracted increasing attention for its biotechnological potential, particularly in the context of plant growth promotion. Several strains act as plant growth-promoting bacteria (PGPB) through mechanisms such as phosphate solubilization, nitrogen fixation, phytohormone production, and siderophore synthesis (Lv et al., 2022; Nascimento et al., 2020; Quecine et al., 2012). In addition, they can enhance plant resilience under stress, including mitigating drought-induced water stress (Luziatelli et al., 2020). The genus biotechnological potential also extends to its application as a bioinoculant and biocontrol agent, antagonizing phytopathogens and stimulating plant innate immunity (Duchateau et al., 2024). These characteristics underscore the genus potential for sustainable crop management.

Despite this promise, the genus also harbors pathogenic species. *Pantoea stewartii*, for example, causes Stewart’s wilt in sweet corn (Ibrahim et al., 2022), while *Pantoea ananatis* is responsible for rice grain discoloration (Doni et al., 2021). Pathogenic strains of *Pantoea agglomerans* induce gall formation in gypsophila and beet (Lorenzi et al., 2022), and additional cases of pathogenic isolates have been reported from other plants such as onion and eucalyptus (Duan et al., 2025; Vahling-Armstrong et al., 2016).

Beyond plants, *Pantoea* can occasionally infect humans, although such cases are much rarer and less studied. Reported clinical manifestations include nosocomial pneumonia, urinary tract infections, bacteremia, and wound infections (Ruan; Qin; Li, 2022; Wang; Liang; Hu, 2024).

This duality—beneficial versus pathogenic lifestyles—poses a critical challenge for the safe application of *Pantoea* in biotechnology. Phenotypic plasticity complicates the screening of strains for safe use as bioinoculants, a challenge shared with other bacterial genera (Tariq et al., 2022). Moreover, despite advances in understanding the mechanisms of phytopathogenicity, the genetic basis underlying the diversity of lifestyles within the genus remains incompletely characterized.

In this study, we address this knowledge gap using comparative genomics of the *Pantoea* genus. We identify species with the greatest biotechnological potential, particularly as biofertilizers, and assess their safety by examining genetic determinants associated with phytopathogenic and clinical lifestyles.

## Methods

### Genome curation and reclassification

A total of 1,286 *Pantoea* spp. genomes available in the GenBank database (accessed February 10, 2025) were retrieved. Genome quality was assessed using CheckM v1.0.13 (Parks et al., 2015), applying a minimum completeness threshold of 90% and a maximum contamination of 10%. Genomes containing more than 500 contigs were discarded. To remove redundancy, in-house scripts based on Mash v2.3 (Ondov et al., 2016) were used to cluster genomes with a Mash distance below 0.05 (∼99.95% ANI) (Passarelli-Araujo; Franco; Venancio, 2022). Within each cluster, the genome with the highest N50 was retained as the representative. Subsequently, ANI comparisons were then performed using ANIm (MUMmer-based alignment) implemented in pyANI v0.2.7 (Pritchard et al., 2016).

### Gene prediction, mobile elements, and phylogeny

Protein-coding sequences were predicted using Prokka v1.13.4 (Seemann, 2014). Mobile genetic elements were identified through plasmid prediction with PlasForest v1.4 (Pradier et al., 2021) and genomic island detection with IslandViewer 4 (http://www.pathogenomics.sfu.ca/islandviewer/, accessed March 27, 2025) (Bertelli et al., 2017). Core single-copy orthologs were identified using OrthoFinder v3.1.0 (Emms; Kelly, 2019). These orthologs were aligned with MAFFT (Katoh; Standley, 2013) and used to reconstruct a maximum likelihood phylogeny with IQ-TREE v2.4.0 (Nguyen et al., 2015). The best-fitting model (Q.plant+F+R10) was determined with ModelFinder, and clade support was assessed with 1,000 bootstrap replicates. Phylogenetic trees were visualized using iTOL v7.2.1 (Letunic; Bork, 2019).

### Super-pangenome analysis and Pan-GWAS

The super-pangenome of the genus *Pantoea* was reconstructed using Roary v3.13.0 (Page et al., 2015), applying an 80% identity threshold for ortholog clustering. A genome-wide association study (GWAS) was then conducted with Scoary v1.6.16 (Brynildsrud et al., 2016), using lifestyle categories (e.g., beneficial, phytopathogenic, clinical) as phenotypes. Input data included the gene presence/absence matrix from Roary and the previously generated phylogenetic tree to account for population structure. To reduce phylogenetic bias, Scoary’s permutation test was applied with 1,000 randomizations of lifestyle labels. Statistical significance was determined by adjusting p-values with the Benjamini–Hochberg method. Genes were considered significant when meeting three criteria: specificity > 90%, adjusted p ≤ 0.05, and empirical p ≤ 0.05.

### Mining of genes of interest

Genes functionally related to host interaction were identified using Usearch v11.0.667 (Zhou; Liu; Li, 2024), with minimum thresholds of 50% identity and 80% coverage. Searches were performed sequentially against: 1) an in-house curated database of plant growth-promoting genes; 2) the Virulence Factors of Pathogenic Bacteria Database (VFDB) (Liu et al., 2022); 3) the Pathogen Host Interaction database (PHI-base) (Urban et al., 2025); and 4) the Comprehensive Antibiotic Resistance Database (CARD) (Alcock et al., 2023). The latter three databases were used to identify virulence- and resistance-associated genes, representing the virulome and resistome of the genus. All databases were accessed in May 2025.

## Results and Discussion

### Genome filtering, species delimitation, and phylogeny

To enable robust comparative analyses and genetic mining, we first curated the genomic dataset, ensuring quality, removing redundancy, and clarifying taxonomic assignments. In total, 1,286 genomes of *Pantoea* spp. were retrieved from the NCBI GenBank database (February 2025). From this initial set, we selected genomes containing fewer than 500 contigs. We then assessed assembly completeness by calculating the percentage of expected single-copy genes, retaining only those with >90% completeness. In addition, we screened for multiple copies of single-copy genes and excluded genomes with >10% contamination. After applying these quality filters, 1,151 genomes were retained.

Redundancy was reduced using Mash, which clustered genomes with pairwise distances below 0.05 (corresponding to 99.95% ANI), identifying 390 duplicate assemblies. Within each cluster, the assembly with the highest contiguity (N50) was selected as the representative genome. This process resulted in a final dataset of 645 high-quality, non-redundant genomes that were used for subsequent analyses, Table S1 lists all downloaded genomes, indicating duplicates and reasons for exclusion.

Species delimitation was performed using an all-vs-all ANI analysis. Genomes sharing ≥ 95% ANI (Bobay, 2020), the accepted threshold for bacterial species boundaries, were grouped together. This analysis identified 189 necessary reclassifications, the majority (80.4%) corresponding to previously unclassified strains. Interestingly, some discrepancies involved type strains themselvess—for example, the type strain of *P. leporis* shares 98.5% ANI with *P. endophytica*, which also possesses its own type strain. Additional cases of misassignment are provided in Table S2.

A maximum likelihood phylogeny was then reconstructed using single-copy orthologs genes identified by OrthoFinder (see Methods for details). The reconstruction resolved 23 major groups, each containing a type strain. These phylogenetic clusters were congruent with the ANI-based species boundaries, reinforcing the robustness of both approaches for taxonomic delimitation within the genus *Pantoea* (Figure 1).

**Figure 1.**
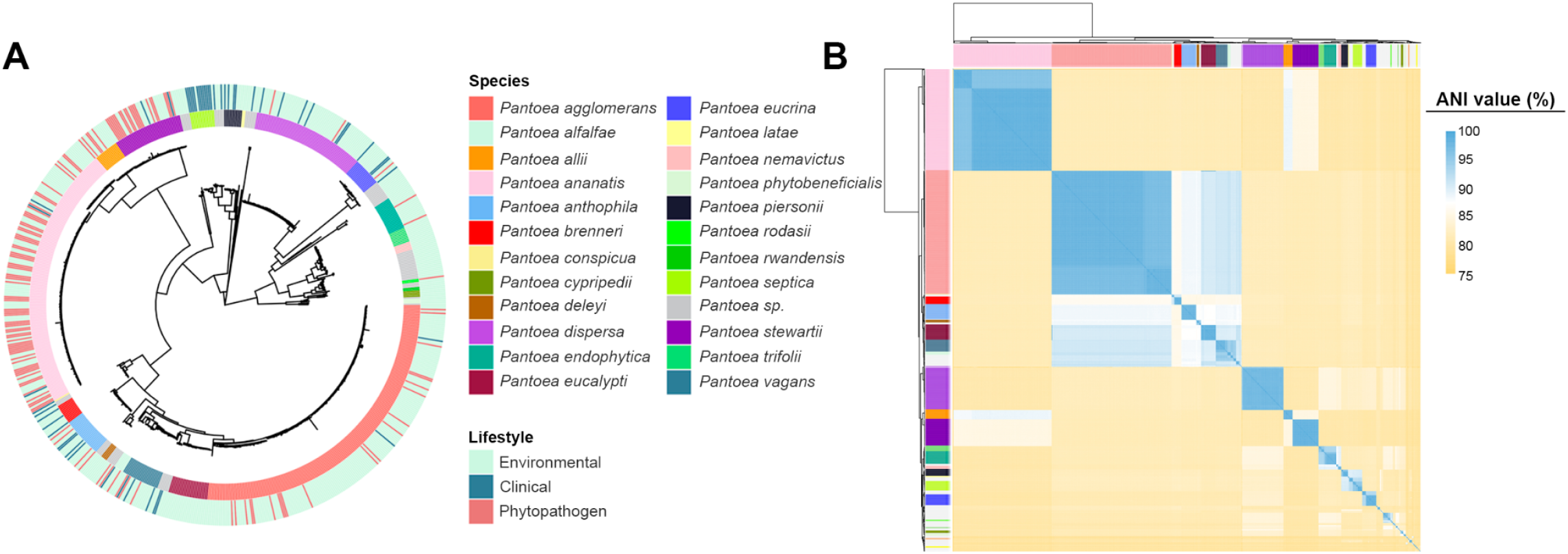
Phylogeny and diversity of the *Pantoea* genus. (A) Maximum likelihood phylogenetic tree of *Pantoea* genomes, indicating species and their associated lifestyles. Reconstruction was performed with IQ-TREE. (B) ANI-based clustering showing 23 genomic groups defined by >95% identity.

Species distribution across lifestyles revealed distinct trends. The most frequently represented phytopathogenic taxa were *P. ananatis, P. stewartii, P. allii*, and *P. agglomerans*, consistent with their recurrent association with crop diseases. In contrast, strains isolated from clinical settings were predominantly assigned to *P. septica, P. piersonii, P. brenneri*, and *P. anthophila*. The remaining species were mostly of environmental origin, commonly associated with plants, and lacked records of pathogenicity or deleterious interactions (Figure 1).

### Plant growth-promoting potential of *Pantoea*

After defining genomic groups and reclassifying the genomes, we examined the repertoire of genes with potential biotechnological relevance, focusing on those associated with plant growth promotion. Strains of *Pantoea* have been increasingly recognized as candidates in agricultural biotechnology, given their documented effects on plant development (Walterson & Stavrinides, 2015). To explore this potential, we screened the genus for classical PGPB traits, including genes involved in nitrogen fixation, phosphate solubilization, phytohormone production, siderophore synthesis, and ethylene modulation. The genes used for genome mining and their predicted functions are listed in Table S3.

Figures 2 and 3 show presence–absence profiles of PGPB-related genes, revealing a broadly homogeneous distribution across *Pantoea* lifestyles. Notably, even phytopathogenic and clinical strains harbor genes traditionally linked to plant growth promotion. These findings indicate that the mere presence of such genes is insufficient to distinguish beneficial from pathogenic lifestyles within the genus. A more detailed discussion of the principal PGPB-related genes identified is provided below.

**Figure 2.**
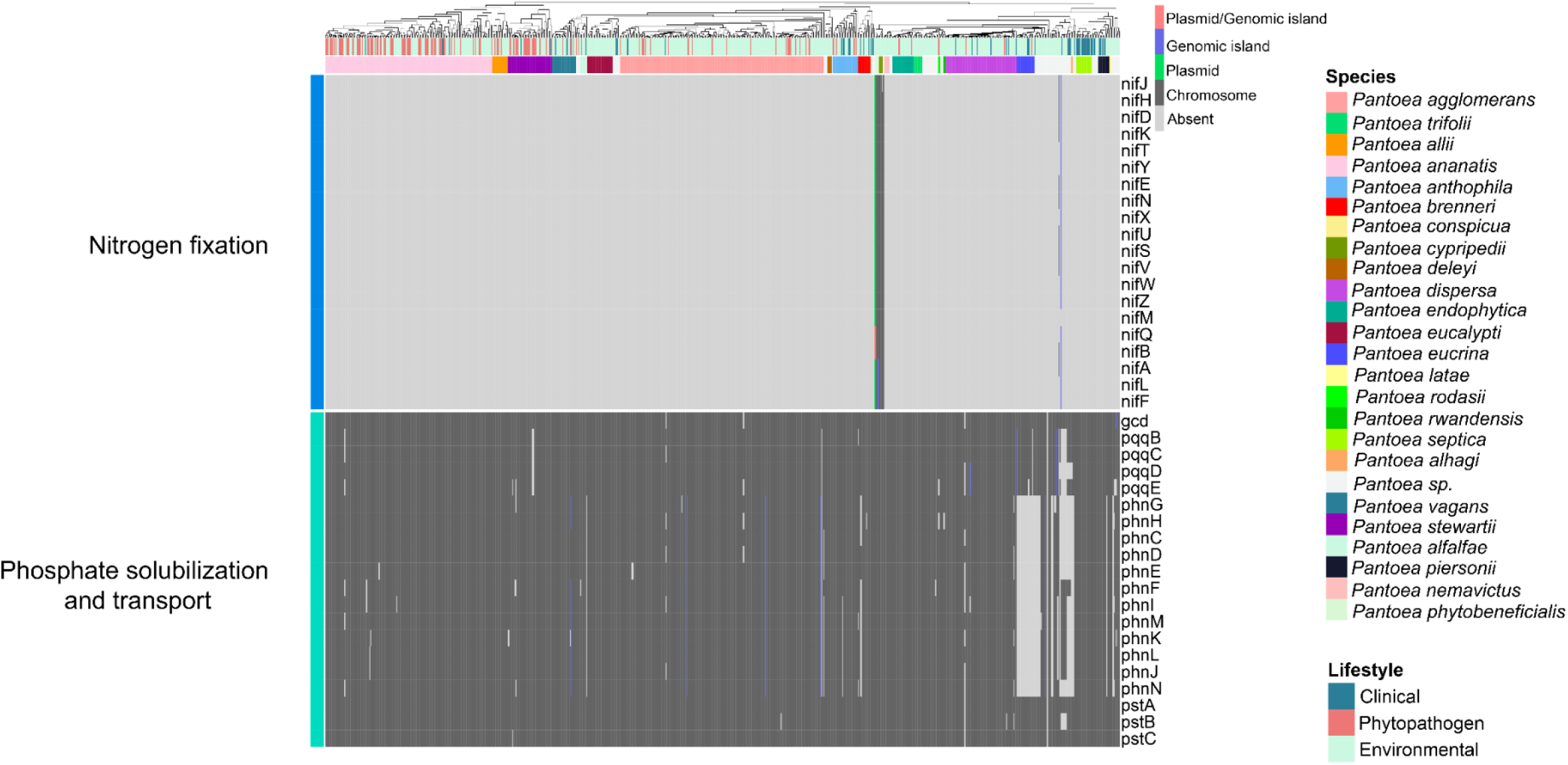
Presence–absence heatmap of genes associated with nitrogen fixation, phosphate solubilization and transport. Columns correspond to individual genomes and are annotated with species names and their respective lifestyles. The occurrence of the same gene in different genomic regions is represented by distinct colors.

**Figure 3.**
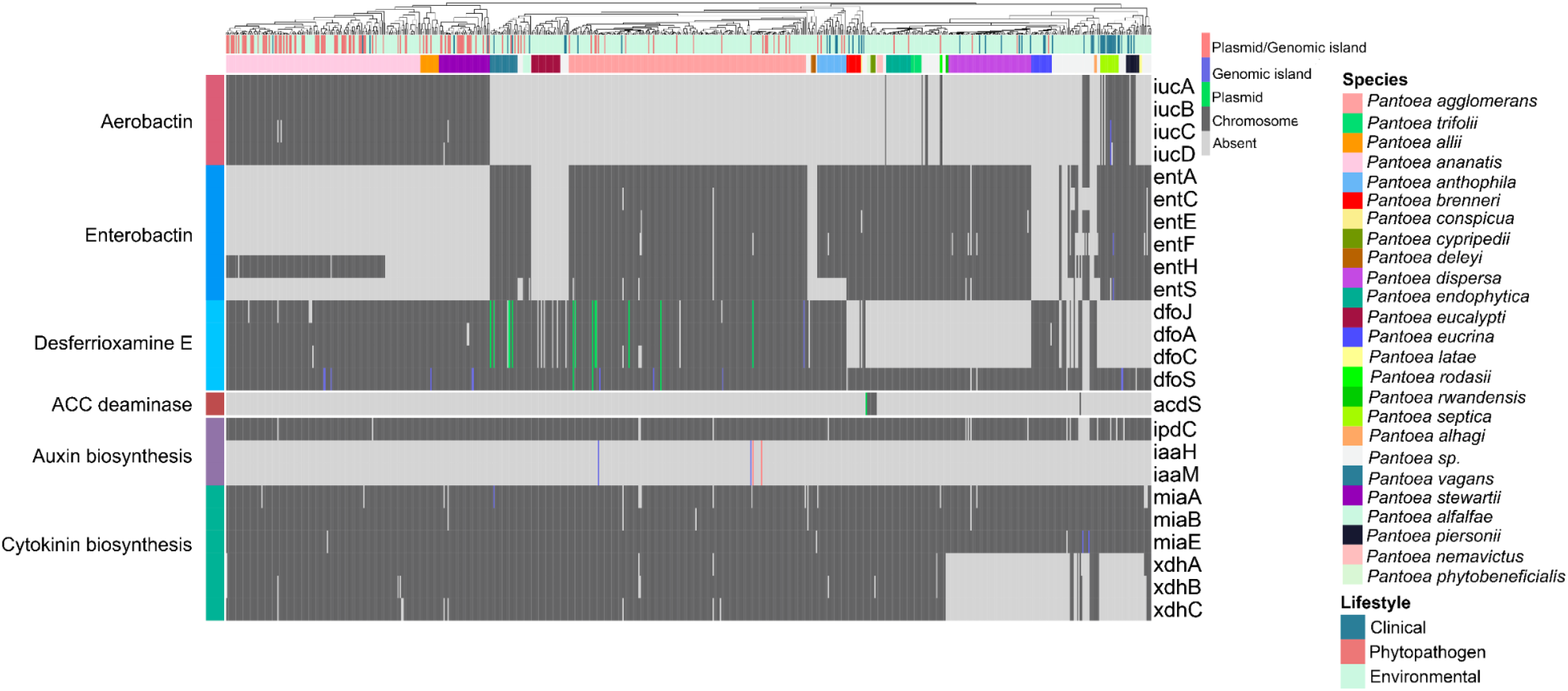
Presence–absence heatmap of genes associated with siderophore biosynthesis, phytohormone production, and ethylene level modulation. Columns represent individual genomes and are annotated with species and their respective lifestyles. Distinct colors indicate the occurrence of the same gene in different genomic regions.

#### Nitrogen fixation and phosphate solubilization

A classical trait of PGPB is the ability to fix atmospheric nitrogen, converting it into forms readily assimilable by plants (Sepp et al., 2023). This function is mediated by the *nif* operon, which encodes approximately 20 proteins involved in nitrogen uptake, biosynthesis of the iron–molybdenum cofactor, reduction of nitrogen to ammonia, and regulatory functions.

All *nif* genes were detected exclusively in *P. cypripedii, P. phytobeneficialis*, and two unclassified strains, forming a phylogenetically close group (Figure 2). In one of these strains, the *nif* cluster is located on a plasmid that also harbors the genes *acdS* and *acdR* (associated with ACC deaminase activity) (Figure 3), as well as genes related to iron transport, jasmonic acid biosynthesis, and glucose-1-dehydrogenase. In *P. cypripedii* and *P. phytobeneficialis*, the *nif* genes are chromosomally encoded and syntenic, suggesting integration into the genome after plasmid acquisition through a horizontal transfer event (HGT). Thus, nitrogen fixation in *Pantoea* is circumscript to a small monophyletic group that likely acquired such genes through a common HGT event.

Phosphate solubilization represents another crucial PGPB trait, given the limited availability of plant-assimilable phosphorus in most soils (Olanrewaju; Glick; Babalola, 2017). Genes related to inorganic phosphate solubilization, such as *gcd* (glucose-1-dehydrogenase) and its cofactor pqq (pyrroloquinoline quinone), which mediate gluconic acid (GA) synthesis, were identified. GA promotes soil acidification, thereby releasing phosphate ions for plant uptake. Genes associated with GA production were consistently detected in chromosomes of all *Pantoea* species, suggesting that phosphate solubilization is an intrinsic feature of the genus.

In addition, the *phn* operon, which encodes phosphonatase—an enzyme responsible for the degradation of organophosphonate compounds that can also serve as a phosphorus sources—was detected in the chromosomes of most species. Notable exceptions included *P. eucrina, P. coffeiphila*, and several unclassified strains.

#### Stress control, phytohormone production, and siderophore biosynthesis

Under biotic and abiotic stress conditions, plants tend to accumulate ethylene, a key hormone that regulates various physiological processes but, at high levels, can inhibit root development and root hair formation (Ahemad; Kibret, 2014). Certain endophytic bacteria mitigate these effects by producing 1-aminocyclopropane-1-carboxylate (ACC) deaminase, which metabolizes ACC, the direct precursor of ethylene in the Yang cycle. The gene encoding this enzyme, *acdS*, was identified in only two species, *P. cypripedii* and *P. phytobeneficialis* (Figure 3). As observed for nitrogen fixation genes, the restricted distribution of *acdS* suggests a high degree of specialization for beneficial plant interactions, reinforcing the potential of these species as biofertilizers.

Auxin, particularly indole-3-acetic acid (IAA), is another phytohormone central to plant development, especially in root growth and cell differentiation (Tang et al., 2023). Several bacterial pathways synthesize IAA, named according to their intermediates. In *Pantoea*, the predominant pathway is the indole-3-pyruvic acid (IPyA) route, catalyzed by indole-3-pyruvate decarboxylase, encoded by *ipdC*. This gene was detected in all species, located on chromosomes, indicating an ancestral and conserved origin. In contrast, the indole-3-acetamide (IAM) pathway, dependent on *iaaM* and *iaaH*, which encode tryptophan monooxygenase and indole-3-acetamide hydrolase, respectively, was found only in four *P. agglomerans* strains, including the pathogenic pathovars *P. agglomerans* pv. *betae* and pv. *gypsophila*, both associated with gall formation. In these strains, IAM genes occur within genomic islands and are often linked to virulence determinants such as the *hrp/hrc* operon, which encodes components of the type III secretion system (T3SS) and the induction of tumor formation (Barash; Manulis-Sasson, 2009). These findings are consistent with previous reports showing that the IAM pathway is characteristic of phytopathogens (Ahemad; Kibret, 2014), whereas the IPyA route has a broader distribution across environmental, pathogenic, and clinical strains, making its role less specific to lifestyle.

In addition to auxin, genes associated with the production of other phytohormones, such as cytokinin—a key regulator of cell division, seed germination, and seed development—were also identified (Orozco-Mosqueda; Santoyo; Glick, 2023). This pathway involves genes related to tRNA modification and recycling (*miaA, miaB*, and *miaE*), as well as subunits of xanthine dehydrogenase (*xdhA, xdhB*, and *xdhC*) (Rocha et al., 2023). Some species, including *P. dispersa, P. eucrina, P. latae, P. septica*, and *P. piersonii*, lacked the *xdhABC*. However, these species harbor *yagTSR*, which also encodes a xanthine dehydrogenase (Neumann et al., 2009), enabling cytokinin biosynthesis (Hossain, 2024).

Siderophores, iron-chelating molecules produced by rhizospheric and endophytic bacteria, enhance iron availability to plants and contribute to pathogen suppression by outcompeting fungal siderophores (Olanrewaju; Glick; Babalola, 2017). Three siderophore types were detected in *Pantoea*: enterobactin (*ent* operon), desferrioxamine E (DFO-E, *dfo*), and aerobactin (*iuc*). While enterobactin and DFO-E-related genes are broadly distributed with occasional losses and no clear lifestyle correlation, aerobactin genes show a more restricted pattern, occurring mainly in *P. ananatis, P. allii*, and *P. stewartii* (primarily phytopathogens), and *P. septica* (a clinical species). These results support previous observations that an enterobactin-like cluster was present in the last common ancestor of the genus, with subsequent losses in certain species and independent acquisitions of other siderophore clusters (e.g. aerobactin), via HGT (Soutar; Stavrinides, 2018).

In enterobacterial human and animal pathogens, aerobactin is a recognized virulence factor, essential for infection and host survival, with decisive roles demonstrated in pathogenic *Escherichia coli* (Pakbin; Brück; Rossen, 2021) and hypervirulent *Klebsiella pneumoniae* (Lim et al., 2025). In contrast, plant-associated bacteria often employ aerobactin in niche colonization and competitive interactions rather than pathogenicity (Choi et al., 2022). Although our pan-GWAS did not reveal significant associations between aerobactin genes and pathogenic lifestyles, their distribution suggests that acquisition of these clusters has conferred adaptive advantages, facilitating colonization of new ecological niches and hosts.

### Antibiotic resistance potential

For the safe implementation of *Pantoea* strains with high biotechnological potential, it is crucial to assess their resistome, i.e., the set of genes associated with antibiotic resistance, in order to avoid establishing genetic reservoirs that could facilitate the spread of clinically relevant resistance determinants (Larsson; Flach, 2022). Our analysis revealed that the two predominant resistance mechanisms in the genus are efflux pump-mediated export and antibiotic inactivation by enzymes.

The distribution of resistance genes does not follow a lifestyle-specific pattern (Figure 4), with clinical strains not showing a distinct repertoire that could discriminate them from environmental, endophytic, or other isolates. It is noteworthy that although some resistance genes were associated with mobile elements, no resistance islands carrying genes related to last-generation antibiotics, such as *bla*_CTX-M-15_ (third-generation cephalosporins) (Darby et al., 2023), *bla*_OXA-48-_like (carbapenems) (Hendrickx et al., 2021), and *qnr* (fluoroquinolones) (Amereh et al., 2023), were observed.

**Figure 4.**
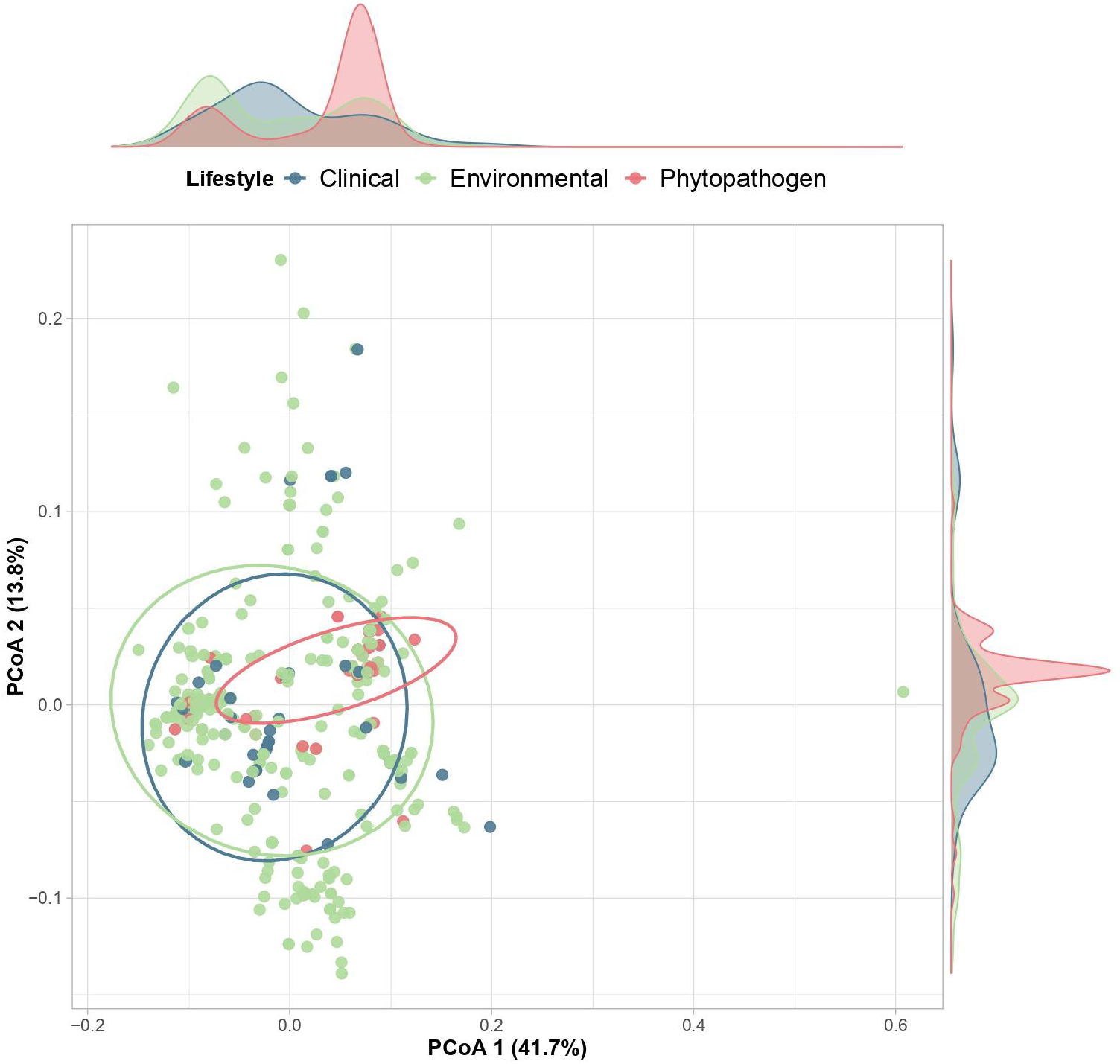
Principal Coordinate Analysis (PCoA) of dissimilarities in antibiotic resistance gene content across *Pantoea* genomes with different lifestyles. Density plots along the axes illustrate the distribution of genomes relative to each coordinate.

This suggests that the *Pantoea* resistome does not reflect recent adaptations driven by strong selective pressures in clinical environments, as seen in bacteria involved in hospital outbreaks (Sanikhani et al., 2024). Thus, the resistance repertoire found appears to represent intrinsic genetic characteristics of the genus rather than recent acquisitions of new mechanisms against modern antibiotics.

#### Efflux pumps and antibiotic inactivation

Efflux pumps are key bacterial defense systems that expel toxic compounds, including antibiotics, thereby enhancing survival in adverse environments (Huang et al., 2022). A wide diversity of efflux pump genes is distributed across the chromosomes of the analyzed strains, many of which are linked to the export of multiple antibiotic classes (Figure 5). This pattern indicates that resistance mechanisms are intrinsic to the genus.

**Figure 5.**
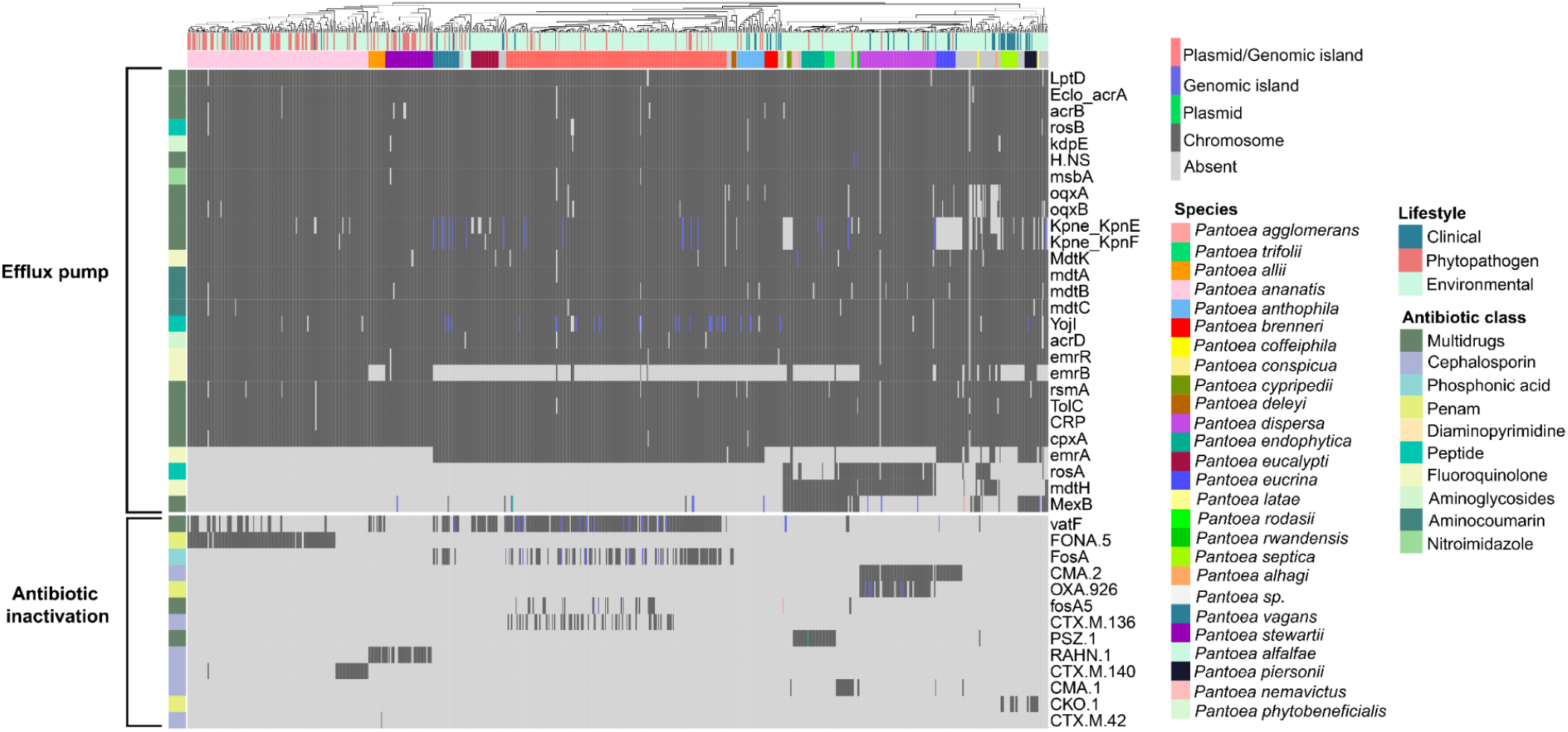
Presence–absence heatmap of genes related to efflux pumps and antibiotic inactivation. Genes are grouped according to the antibiotic classes to which they confer resistance, while genomes are labeled by species and lifestyle.

Among the efflux pump superfamilies identified, the most prominent were RND (Resistance-Nodulation-Division), MFS (Major Facilitator Superfamily), and SMR (Small Multidrug Resistance). The *acrAB* and *oqxAB* complexes, belonging to the RND superfamily, mediate the extrusion of several antibiotics, particularly cephalosporins and fluoroquinolones. These systems confer resistance to second- and third-generation fluoroquinolones as well as third- and fourth-generation cephalosporins in clinical pathogens such as *E. coli* and *K. pneumoniae* (Bialek-Davenet et al., 2015; Huguet; Pensec; Soumet, 2013). Additional relevant genes were also detected, including *rosAB* (MFS), associated with resistance to antimicrobial peptides (Bengoechea; Skurnik, 2000); *kpnEF* (SMR), linked to resistance to cephalosporins and tetracyclines (Srinivasan; Rajamohan, 2013) ; *mdtABC* (RND) (Nishino; Nikaido; Yamaguchi, 2007), involved in resistance to aminocoumarins; and *emrB-tolC* (MFS), associated with fluoroquinolone resistance (Gu et al., 2021).

The widespread presence of diverse efflux pumps across the genus is common in soil and rhizosphere bacteria (Delgado-Baquerizo et al., 2022). Beyond their roles in antibiotic resistance, these efflux pumps contribute to physiological processes including microbial competition and cell-to-cell signaling, with antibiotic resistance representing an exaptation of these functions (Blanco et al., 2016).

With respect to enzymatic antibiotic inactivation, 13 genes were identified, but their distribution was restricted to specific species. Most of these genes correspond to β-lactamases active against cephalosporins, found in *P. ananatis, P. stewartii, P. allii, P. agglomerans, P. dispersa*, and *P. eucrina*. Additional clinically relevant enzymes included *fosA*, conferring resistance to fosfomycin, present in *P. vagans* and *P. agglomerans*, and *PSZ-1*, a β-lactamase originally reported in *P. endophytica* and here confirmed to be present in all genomes of this species, as well as its sister species, *P. trifolii*.

While genomic determinants suggest a potential for clinically relevant antibiotic resistance, phenotypic susceptibility testing of clinical *Pantoea* isolates has shown broad susceptibility to the majority of antibiotics tested, with resistance primarily observed to ampicillin, fosfomycin, and piperacillin/tazobactam (Casale et al., 2023).

### Genetic determinants of *Pantoea* lifestyles

The genus *Pantoea* is well known for its ecological versatility, comprising strains with diverse lifestyles, including environmental, plant-associated, and even animal and human-associated forms. In this study, we classified the analyzed strains into three main groups: environmental, phytopathogenic, and clinical. Within the environmental group, endophytic species are particularly noteworthy, as they colonize plant tissues without causing visible symptoms (Hallmann et al., 1997). However, the boundary between endophytic and phytopathogenic bacteria is often blurred, since both share similar genetic repertoires, especially virulence factors, involved in plant colonization and infection (Lòpez-Fernàndez et al., 2015). This genetic overlap complicates the identification of truly beneficial strains and represents a key challenge when screening microorganisms for biotechnological applications.

To further explore these distinctions, we performed a pan-GWAS analysis to identify accessory genes associated with the different lifestyles and pinpoint genetic determinants that could serve as discriminators among the categories. Genes significantly associated with the phytopathogenic lifestyle (see Methods for details) included components of the T3SS, specific T3SS effectors, and the central gene of the phosphonate biosynthetic cluster (HiVir). The restricted distribution of these determinants among phytopathogenic strains is shown in Figure 6 and Figure 7.

**Figure 6.**
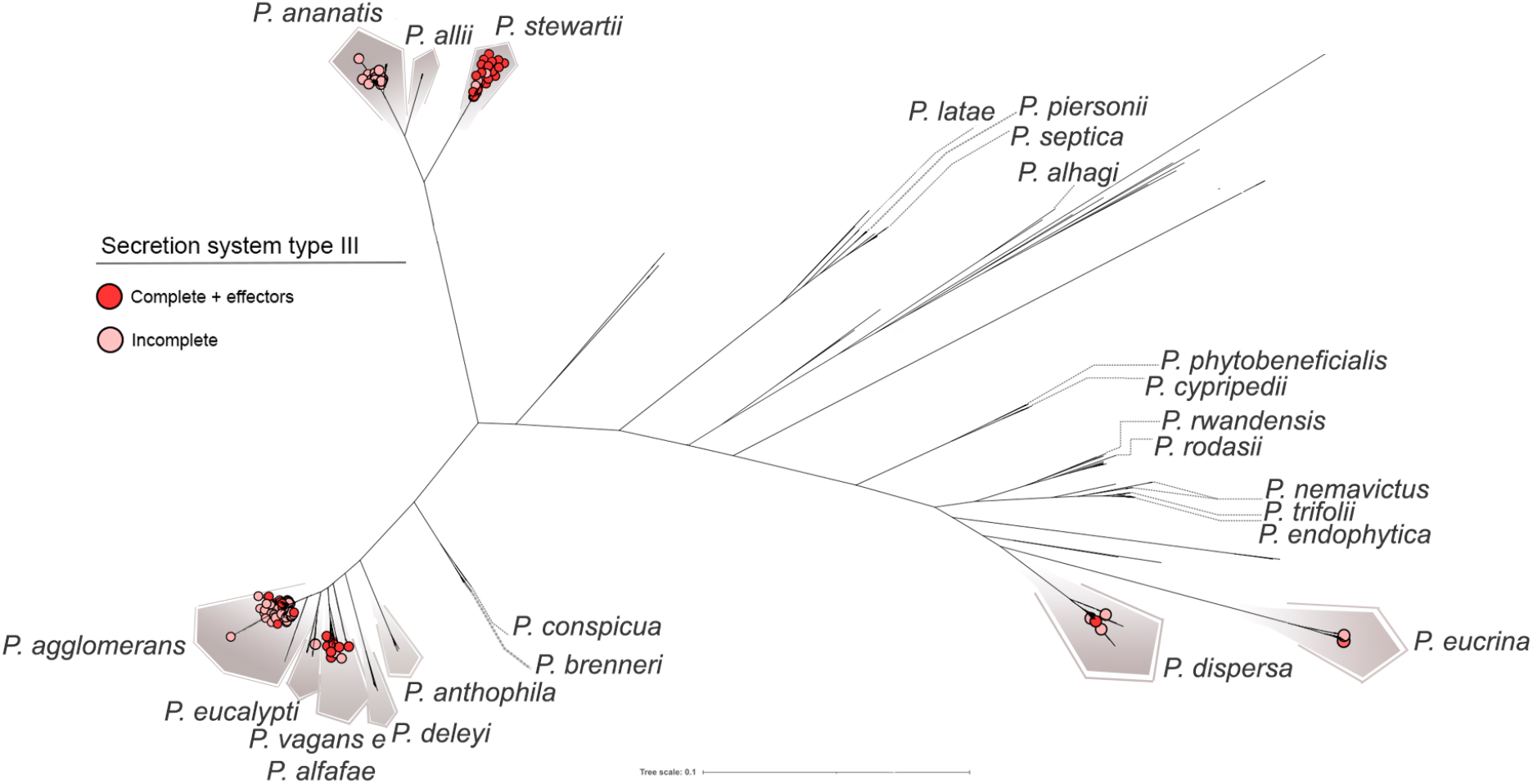
Distribution of the type III secretion system (T3SS) of the *hrp-hrc* family across *Pantoea* species. Dark red circles denote strains carrying the complete *hrp-hrc* cluster together with its effectors, while light red circles represent strains lacking the full cluster and associated effectors. The complete system is mainly found in *P. stewartii, P. agglomerans, P. vagans*, and *P. alfalfae*, which harbor all genes encoding the T3SS apparatus as well as the effectors linked to host specificity and pathogenicity—for example, *wtsE* in *P. stewartii* (water-soaking symptoms), and *hsvB* and *hsvG* (host specificity and gall formation). The restricted distribution of the T3SS among phytopathogenic species suggests its potential as a genetic marker to distinguish pathogenic from beneficial *Pantoea* strains.

**Figure 7.**
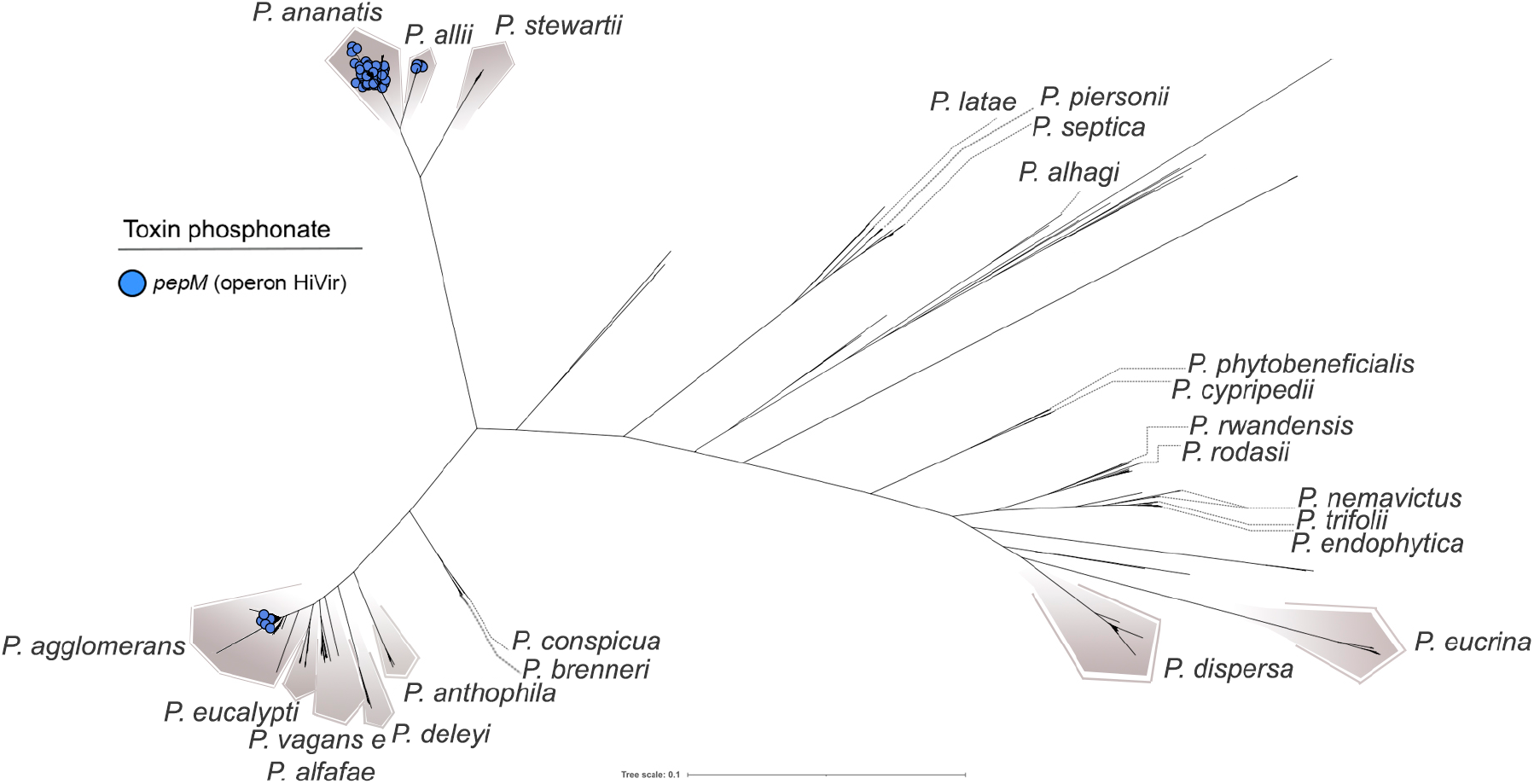
Distribution of the *pepM* gene across species of the genus *Pantoea*. The *pepM* gene is part of the HiVir operon, which encodes enzymes for the biosynthesis of pantaphos, a phosphonate phytotoxin implicated in plant disease. Species harboring *pepM* are highlighted, illustrating its restricted distribution and potential association with virulence and host specificity.

**Figure 8.**
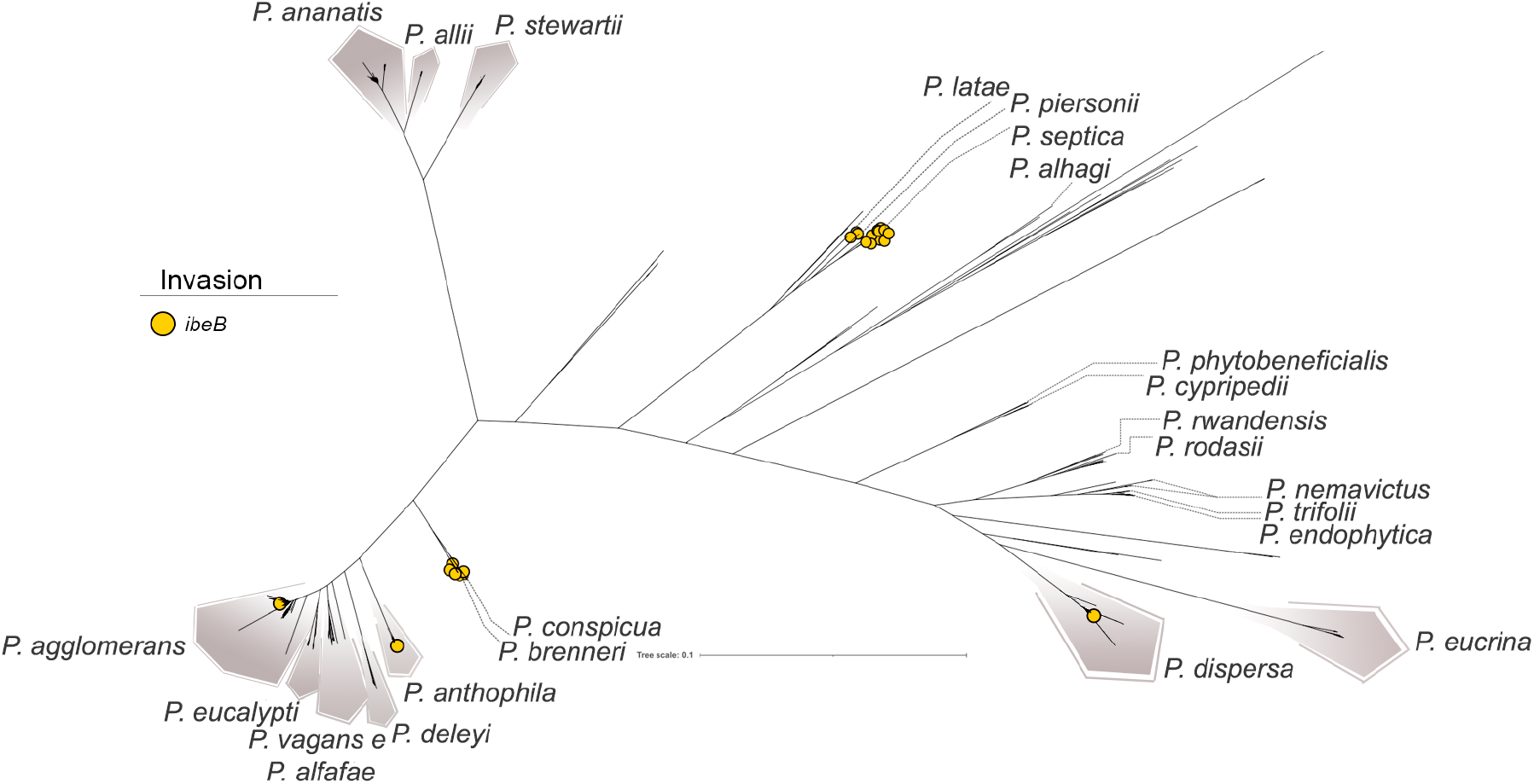
Distribution of the *ibeB* gene across *Pantoea* species, highlighting its prevalence in strains associated with clinical or pathogenic lifestyles. The presence of *ibeB* suggests a potential role in invasiveness and may serve as a marker for distinguishing pathogenic from environmental isolates.

A key determinant of pathogenicity in Gram-negative phytopathogens is the T3SS, a membrane-embedded nanomachine composed of ∼20 structural proteins. The T3SS is encoded by the *hrp/hrc* gene cluster, whose distribution appears to be restricted to a limited number of species (Figure 6). This system mediates the translocation of effector proteins into plant host cells through a needle-like pilus structure, known as the Hrp pilus. Once inside the host cells, these effectors can either trigger plant immune responses or suppress host defenses to promote disease development (He et al., 2025; Ji; Dong, 2015).

Among the analyzed species, *P. stewartii* harbored the complete *hrp/hrc* cluster in most genomes. The cluster was also detected in five *P. vagans* strains (plasmid-borne in one of them) and in the pathovars *P. agglomerans* pv. *gypsophila* and pv. *betae. P. stewartii* subsp. *stewartii* is a well-known pathogen causing Stewart’s bacterial wilt and leaf blight in maize (*Zea mays* L.), with the *hrp/hrc* cluster being essential for its pathogenicity (Mergaert; Verdonck; Kersters, 1993). In *P. agglomerans*, these pathovars are established pathogens of *Gypsophila* and sugar beet, respectively.

Consistent with observations on phytohormone production, the *hrp/hrc* operon is also associated with gall formation, acting synergistically with auxin biosynthesis genes of the IAM pathway (*iaaM* and *iaaH*). In several instances, these determinants were plasmid-encoded, suggesting the horizontal acquisition of pathogenicity islands that enabled typically endophytic strains to adopt a pathogenic lifestyle.

Among the effectors, *wtsE* stands out as essential for lesion development in maize by *P. stewartii*, where it induces water-soaking and facilitates bacterial proliferation (Jin et al., 2016). This effector was also detected in *P. agglomerans* pv. *gypsophila* and pv. *betae*. Along with this, the effectors *hsvG* and *hsvB* were exclusively detected in these pathovars, consistent with their specialized interactions with *Gypsophila* and sugar beet.

Recent reports have also described plant lesions caused by *P. vagans* (Anwer et al., 2025; Rodríguez Velázquez et al., 2024), a species characterized by diverse lifestyles. Some of its strains, such as *P. vagans* C9-1, are recognized for their biotechnological applications and use as biocontrol agents (Klein et al., 2017), further highlighting the dual ecological roles of this species.

Overall, the T3SS emerges as a central determinant of pathogenicity in different *Pantoea* strains infecting diverse plant hosts. Its recurrent occurrence underscores its potential as a promising target for novel control strategies, including molecular inhibition of T3SS components, an approach already supported by recent studies (Anwer et al., 2025).

However, not all *Pantoea* phytopathogens rely on the T3SS. A notable exception is *P. ananatis*, widely studied for its role in onion center rot (Gitaitis et al., 2002; Shin; Dutta; Kvitko, 2023; Stice et al., 2018). Unlike other pathogenic species in the genus, *P. ananatis* lacks the T3SS and instead depends on the HiVir biosynthetic gene cluster, which encodes the phosphonate phytotoxin pantaphos (Agarwal et al., 2021). Within this cluster, *pepM*, encoding a phosphoenolpyruvate mutase, is essential for pathogenicity in onions (Yang et al., 2025). This phytotoxin is predominantly found in *P. ananatis* (Figure 7), where it is located in genomic islands, suggesting acquisition via HGT. Additionally, *pepM* was identified in four *P. allii* strains (three from onion bulbs showing center rot symptoms), one *P. vagans* strain isolated from asymptomatic *Alliaria petiolata* (Shin; Kvitko, 2024), and seven *P. agglomerans* strains also isolated from diseased onions. In all cases, the genes were located within genomic islands, indicating possible plasmid-mediated transmission of the HiVir cluster across species within the genus.

The pan-GWAS analysis identified a single gene with strong biological plausibility that was significantly associated with clinical strains: *ibeB* (also known as *cusC*), encoding an invasin-related protein. In *E. coli*, ibeB has been described as a virulence factor involved in human neonatal meningitis (Germon et al., 2005), while in *Cronobacter sakazakii*—an opportunistic pathogen responsible for severe neonatal infections such as necrotizing enterocolitis, meningitis, and sepsis—it has been similarly linked to pathogenesis (Kucerova et al., 2010). The *ibeB* gene encodes a component of a silver and copper cation efflux system, and its activity has been associated with bacterial invasiveness. In *E. coli*, for instance, ibeB facilitates the entry of bacteria into brain microvascular endothelial cells, thereby promoting penetration of the blood-brain barrier (Franke et al., 2003). This gene was most frequently detected in *P. septica*, particularly in strains isolated from neonatal feces. This species has been associated with neonatal bloodstream infections, mirroring the pathogenic profiles of *E. coli* and *C. sakazakii*, and suggesting a recurring pattern of neonatal infection. The gene was also identified in other clinically derived *Pantoea* species, including *P. piersonii*, isolated from a bacteremia case (Howard et al., 2023) and kidney stones (Rekha et al., 2020), and *P. dispersa*, associated with bloodstream infections. Notably, *ibeB* was also detected in strains obtained from asymptomatic patients, underscoring its presence beyond overt clinical disease.

The *ibeB* genes were found on plasmids and within genomic islands, suggesting HGT events that may have allowed originally commensal bacteria to acquire the ability to infect humans. While their presence in asymptomatic hosts indicates that *ibeB* alone is not sufficient to cause infection alone, the high frequency of this gene in *P. septica—*particularly in strains associated with neonatal infections—points to a potentially important role in the virulence of this species.

To date, the pathogenic mechanisms of *P. septica* remain poorly characterized, and our data highlight *ibeB* as a candidate genetic determinant linked to human disease, especially in neonates. Other species, including *P. eucrina, P. dispersa, P. ananatis*, and more frequently *P. agglomerans*, have also been reported in clinical contexts. However, the relationship between these species and human disease is still uncertain (Crosby et al., 2023). Our analyses did not identify exclusive genetic markers supporting a direct association with pathogenicity in these groups, suggesting that further studies are warranted to clarify whether such strains act as true opportunistic pathogens or instead represent human commensals.

## Conclusions

In this comparative genomics study, we evaluated the biotechnological potential of the genus *Pantoea* and identified species that appear both promising and safe for practical applications, based on the distinctions among ecological lifestyles. *P. cypripedii* and *P. phytobeneficialis* stand out, as they lack genetic determinants linked to phytopathogenicity or human virulence, while harboring a myriad of key plant growth-promoting traits, including phosphate solubilization, phytohormone production, and iron acquisition. These features underscore their potential as safe bioinoculants. Other species displayed relevant PGPB mechanisms but lacked clear ecological specialization, having been recovered from diverse environments, including insects, urban habitats, and aquatic sources. Some of these species have also been reported in pathogenic or clinical contexts, underscoring the need for further studies to properly balance their beneficial potential versus possible risks.

Within phytopathogenic groups, *P. stewartii* emerges as a genuine pathogen. In contrast, lineages of *P. agglomerans* (including pathovars *betae* and *gypsophilae* and those associated with onion center rot), as well as *P. ananatis* and *P. allii*, represent lineages that acquired phytopathogenic capacity over time. In the clinical context, *P. septica* was particularly notable due to the presence of the *ibeB* gene, a candidate determinant of neonatal sepsis that sheds light on its potential virulence mechanisms.

Taken together, our findings highlight the ecological versatility of the genus *Pantoea*, clarify the functional roles of its major species, and provide a framework for the safe use of selected strains in agricultural biotechnology. At the same time, they point to the need for more detailed investigations into clinical and pathogenic lineages to better delineate risks and opportunities for practical applications.

## Acknowledgements

This work was supported by Fundação Carlos Chagas Filho de Amparo à Pesquisa do Estado do Rio de Janeiro (FAPERJ), Coordenação de Aperfeiçoamento de Pessoal de Nível Superior-Brasil (CAPES; Finance Code 001), Conselho Nacional de Desenvolvimento Científico e Tecnológico (CNPq) and Programa de Apoio à Pesquisa, Inovação e Cultura (PAPIC - UENF). The funding agencies had no role in the design of the study and collection, analysis and interpretation of data and in writing.

## Author contributions

Felipe F Rimes-Casais: Data curation, Formal analysis, Writing – original draft, Writing – review & editing. Francisnei Pedrosa-Silva: Conceptualization, Methodology, Supervision. Thiago M. Venancio: Conceptualization, Funding acquisition, Project administration, Supervision, Writing – original draft, Writing – review & editing.

## Notes

### Competing Interest Statement

The authors have declared no competing interest.

